# scDrugAtlas: an integrative single-cell drug response database for dissecting tumor heterogeneity in therapeutic efficacy

**DOI:** 10.1101/2024.09.05.611403

**Authors:** Yanfei Wu, Wei Huang, Xinda Ren, Hui Liu, Judong Luo

**Affiliations:** College of Computer and Information Engineering, Nanjing Tech University, Nanjing, 211800, Jiangsu, China; Department of Radiotherapy, Tongji Hospital, School of Medicine, Tongji University, Shanghai, 200333, China

**Keywords:** Single-cell transcriptome, Drug response, Drug resistance, Tumor heterogeneity, Pseudotime trajectory

## Abstract

Tumor heterogeneity often leads to substantial differences in responses to same drug treatment. The presence of pre-existing or acquired drug-resistant cell sub-populations within a tumor survive and proliferate, ultimately resulting in tumor relapse and metastasis. The drug resistance is the leading cause of failure in clinical tumor therapy. Therefore, accurate identification of drug-resistant tumor cell subpopulations could greatly facilitate the precision medicine and novel drug development. However, the scarcity of single-cell drug response data significantly hinders the exploration of tumor cell resistance mechanisms and the development of computational predictive methods. In this paper, we propose scDrugAtlas, a comprehensive database devoted to integrating the drug response data at single-cell level. We manually compiled more than 100 datasets containing single-cell drug responses from various public resources. The current version comprises large-scale single-cell transcriptional profiles and drug response labels from 1,023 samples (cell line, mouse, PDX models, patients and bacterium), across 78 unique drugs and 24 major cancer types. Particularly, we assigned a confidence level to each response label based on the tissue source (primary or relapse/metastasis), drug exposure time and drug-induced cell phenotype. We believe scDrugAtlas could greatly facilitate the Bioinformatics community for developing computational models and biologists for identifying drug-resistant tumor cells and underlying molecular mechanism. The scDrugAtlas database is available at: http://drug.hliulab.tech/scDrugAtlas/.

## 1 Introduction

The existence of intrinsic and acquired drug-resistant cells is the main cause of failure in tumor therapy. Despite the fact that routine drug treatment can effectively eliminate the majority of malignant cells, a small subset of tumor cells would survive and continue to proliferate, ultimately leading to tumor recurrence and progression. Some scholars have developed bioinformatic methods to predict patients’ clinical response to drugs based on the bulk RNA-seq of cell lines already with large-scale drug sensitivity data, such as GDSC [1, 2], CCLE [3, 4] and CTRP [5]. However, these bulk-level drug sensitivity data can not identify the response or resistance of individual cells to specific drugs, thereby failing to adequately dissect the tumor heterogeneity in drug responses.

In recent years, the advent of single-cell RNA-sequencing (scRNA-seq) technology had generated a wealth of single-cell transcriptional data, providing valuable opportunity to explore the intratumoral heterogeneity in drug response [6]. However, no large-scale experimental data for single-cell drug sensitivity is available, resulting in a significant gap in our knowledge regarding drug response at the single-cell level and bulk level. This gap press a challenging but urgent task to predict the drug sensitivity of individual cells. Several studies have employed transfer learning techniques to align bulk RNA-seq and single-cell RNA-seq (scRNA-seq) data for predicting drug responses at the single-cell level [7–10]. For example, scDeal [11] integrates bulk RNA-seq and scRNA-seq data by aligning domains via maximum mean discrepancy (MMD) to predict single-cell drug responses. Following this idea, SCAD [12] employs adversarial domain adaptation to predict single-cell drug sensitivity. Moreover, scAdaDrug [13] employs adaptive-weighted multi-source domain adaptation to align scRNA-seq data with multiple bulk RNA-seq data domains, resulting in improved performance in predicting drug responses of individual cells. SSDA4Drug [14], on the other hand, utilizes semi-supervised domain adaptation to enhance domain alignment by leveraging few-shot samples from the target scRNA-seq domain, thereby achieving superior accuracy in identifying individual drug-resistant cells. However, the scarcity of single-cell drug response data still significantly impede the exploration of tumor cell resistance mechanisms, and also restricts the development of high-performance computational methods for predicting drug-resistant individual cells.

So, we have worked hard to develop an integrative single-cell drug response database, referred to as scDrugAtlas. We manually compiled single-cell drug response datasets from various public resources, including GEO, PubMed publications and external databases. To ensure the reliability of the drug response labels, we carefully examined that the cell tissue originated from primary, recurrent, or metastatic lesions, whether the cell line was exposed to the drug for a sufficient duration, and whether the cells acquired resistance and continued to proliferate. Based on these assessments, we assigned response labels to each cell regarding specific drug. After data preprocessing and filtering of low-quality data, our database now comprises the transcriptional profiles and drug response labels of 1,205,843 cells came from more than 1,000 samples (cell line, mouse, PDX models, patients and bacterium), across 78 unique drugs and 24 major cancer types. Furthermore, we have built quite a few integrated datasets for some cancer type of interest by removal of batch effects. To our best knowledge, scDrugAtlas is the largest comprehensive database devoted to single-cell drug response to date.

## 2 Data resources and integration

First, we retrieved the PubMed publications using the keywords “((single cell) OR (scRNA-seq) “ AND “drug” AND “(response) OR (sensitivity) OR (resistance) OR (tolerance)”, and subsequently extracted associated data and determined the response labels of individual cells by manually reviewing the content of these publications. Also, we used the same keywords to retrieve datasets from GEO [15] and determined the single-cell response labels based on the corresponding publications. Finally, we also selected some datasets from external databases, including CeDR [16] and DRMref [17]. Figure 1 illustrated the data resources, preprocessing and integration procedures, as well as data analysis functionalities.

**Fig. 1.**
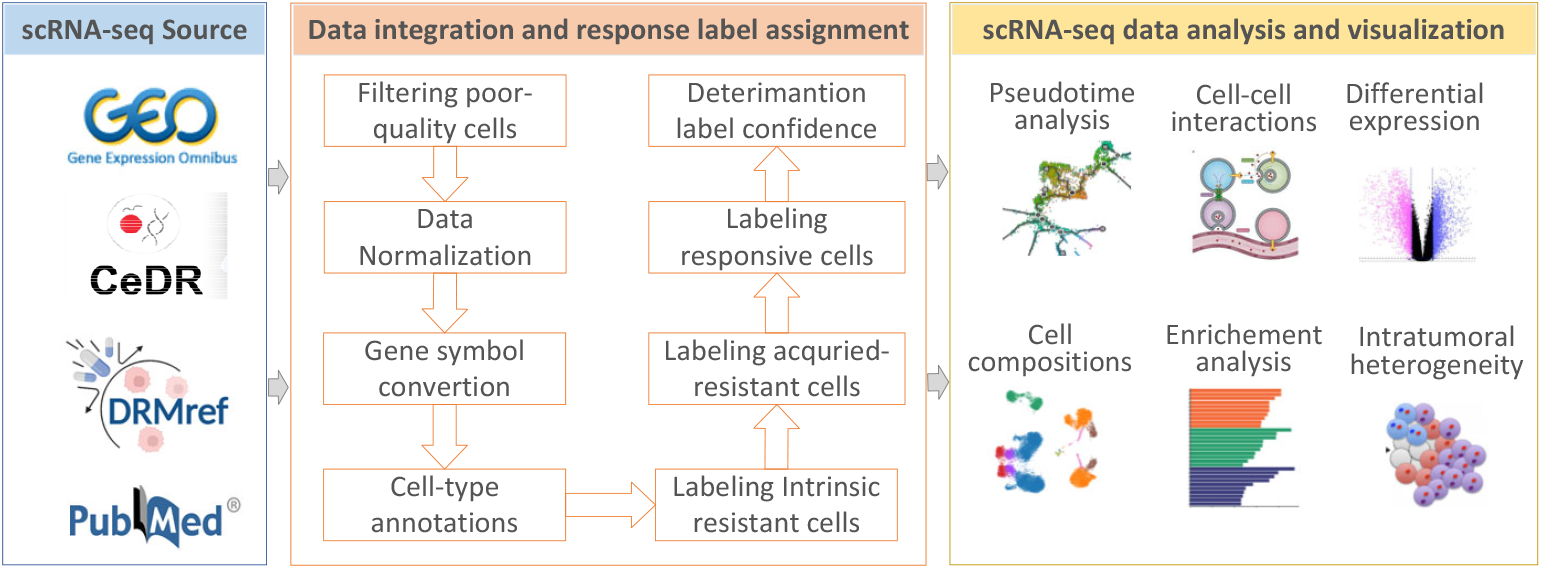
Overview of scDrugAtlas data resource, data integration and drug response label assignment, and functional modules for single-cell transcriptomic data analysis and visualization.

**Fig. 2.**
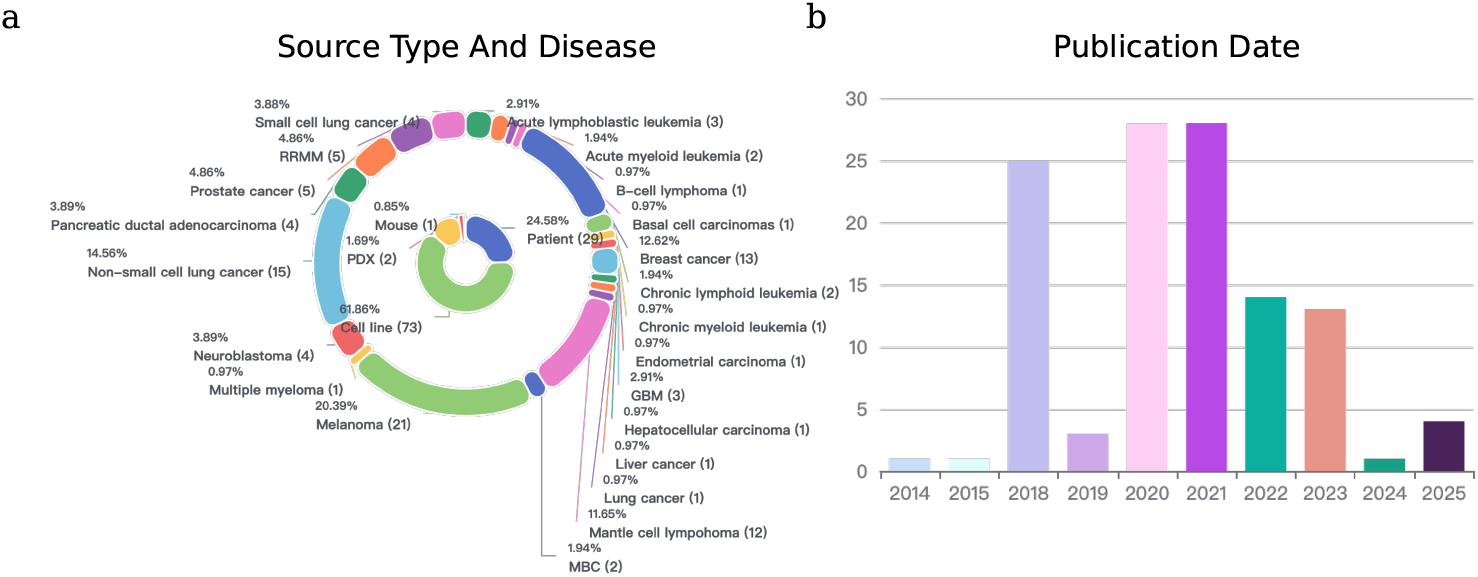
Statistics of scDrugAtlas datasets. (a) Proportion of datasets according to cell source types and diseases. (b) Frequency of datasets according to publication date.

After data collection, we excluded cells with the number of expressed genes *<*500 and with the mitochondrial gene ratio *>*10% from each dataset. Subsequently, we utilized pyensembl package to convert gene identifiers to Ensemble IDs. As a result, scDrugAtlas now comprises the single-cell transcriptional profiles and drug response labels of 1,205,843 cells came from 1,023 samples (cell line, mouse, PDX models, patients and bacterium), across 78 distinct drugs and 24 major cancer types. The drugs covered chemotherapy, targeted therapy and immunotherapy. Detailed drug information was extracted from DrugBank (version 5.1.10) [18].

To facilitate the use of our data resource, we built more than ten integrated datasets for selected cancer type by combining scRNA-seq data from the same cancer type subjected to specific drug treatments. Specifically, datasets were merged, batch labels were assigned, and batch effects were corrected using the batch-balanced kNN algorithm implemented in Scanpy [19]. For example, one integrated dataset comprises 23,314 melanoma cells subjected to Vemurafenib treatment, including 11,955 responsive and 11,359 resistant cells. Another combined dataset consists of 9,274 breast cancer cells treated with Paclitaxel, with 6,235 responsive and 3,039 resistant cells. These integrated datasets can be freely available for download from our database and are ready for use in training machine learning models to predict single-cell drug responses.

## 3 Label assignments and confidence levels

Determining drug-sensitive samples at the single-cell level often presents significant challenges, since single-cell sequencing inherently destroys the physical integrity of the cells. In principle, to determine whether an individual cell responds to a drug requires treating the cell with the drug and observing subsequent changes in its fate. However, once a cell undergoes apoptosis or is effectively killed by drug exposure, it becomes infeasible to conduct single-cell sequencing. As a result, drug-sensitive labels are often assign to the control-group cells based on response of treated groups to drug treatments. Specifically, when cells in the treatment group failed to survive or proliferate after drug exposure, we classified the corresponding control group cells as drug-sensitive.

While untreated control cells were operationally designated as drug-sensitive, those persisting under therapeutic pressure were classified as resistant. To move beyond this binary classification, we developed a systematic pipeline coupled with a confidence-rating scheme to capture both the robustness and stability of resistant phenotypes. This pipeline integrates multiple layers of evidence, including clinical endpoints such as tumor recurrence or metastasis, drug-exposure parameters encompassing treatment duration and biological context, and the proliferative capacity of cells under sustained therapeutic exposure. By jointly considering these criteria, resistant populations were stratified into three tiers of confidence: high, medium, and low. This pipeline facilitates consistent annotation of resistant cell populations and provides a robust foundation for downstream mechanistic studies and the development of predictive models for therapeutic response.

High-confidence resistance is most clearly evidenced by tumor recurrence or metastasis following therapy, as cells derived from these lesions have undergone stringent selective pressure within the complex human microenvironment. Intrinsic resistance likewise reflects stable biological traits that confer constitutive survival advantages. In addition, prolonged drug exposure over extended periods (*>*6 months) has been shown to drive tumor cells toward durable, heritable resistance through genetic mutations, transcriptional reprogramming, or epigenetic remodeling. Robust proliferation under sustained therapeutic pressure therefore represents the hallmark of a fully resistant state. On this basis, cells meeting one or more of the following criteria are classified as high-confidence, three-star resistant: (i) originating from recurrent or metastatic patient samples, PDX models, or mouse tumors; (ii) displaying well-established intrinsic resistance (e.g., NSCLC cells to TKIs such as erlotinib); or (iii) regaining proliferative capacity after more than 6 months of continuous drug exposure.

Medium-confidence resistance corresponds to intermediate drug exposure (1–6 months), which often induces adaptive responses by activating alternative survival pathways that counteract drug-induced apoptosis. Such cells typically occupy a dynamic “drug-tolerant” state, reflecting an intermediate phase in the evolutionary trajectory toward stable resistance. Their limited proliferative potential indicates that resistance mechanisms have not yet fully consolidated. Accordingly, cell populations surviving 1–6 months of treatment but exhibiting limited proliferation are classified as medium-confidence, two-star resistant.

Low-confidence resistance is assigned to cells exposed to short-term treatment (*<*1 month), where stable genetic or epigenetic adaptations are unlikely to emerge. In these cases, survival is more plausibly explained by transient, reversible stress responses rather than true resistance. Comparisons with DMSO-treated controls—which serve as the most reliable sensitive reference—highlight the instability of this phenotype. Cell populations subjected to *<*1 month of exposure are therefore designated as lowconfidence, one-star resistant.

## 4 Functionalities

A website with user-friendly data visualization and analysis tools is provided to help users explore the wealth of data (Fig 3a). Users can input a gene of interest to explore the scRNA-seq data with each cell colored by its drug response label. Other popular analysis tools includes differential expression analysis between sensitive vs resistant cell populations, pseudotime analysis for visualizing the cancer cell progression to drug resistance along with evolutionary trajectory. We summarized the functional modules as below:

- Dataset listing and search: Users can search for single-cell drug response datasets of interest through keywords on the homepage. Each dataset has been associated with tissue or cell source, cell count, drug, and quality level of response labels. In particular, users can filter datasets by setting source type, species, release time, and disease on the Dataset Listing page (Fig 3b).
- Evolutionary trajectory analysis: Pseudotime analysis is performed using monocle3 tool [20]. The visualization of drug sensitivity changes along the evolutionary trajectory of cells is helpful to discover the critical state of tumor cells transitioning from sensitivity to resistance and thus identify key drug-resistant genes (Fig 3c).
- Gene search across datasets: User can search for genes of interest across multiple scRNA-seq datasets and narrow down the scope by species and tissue type. We use UMAP to display the expression levels of the gene across different datasets.
- Differential expression analysis: differential expression analysis between sensitive and resistant cells in a specific dataset or across two datasets can be easily conducted. The volcano plot is used to display differentially expressed genes. Moreover, users can launch functional enrichment analysis using the up-regulated or down-regulated gene sets here.
- Pseudotime analysis: pseudotime analysis is used to visualize the continuous progression of tumor cells along an inferred temporal trajectory (Fig 3d). Each cell is assigned a pseudotime value computed by monocle3 tool, allowing users to capture the gradual transition from early to late cancer cell states, together with the response to specific drug treatments. This module helps in identifying the intermediate states that are most critical during the development of drug resistance, providing valuable insights into the dynamic nature of tumor evolution.
- Functional enrichment analysis: To better explore the biological functions of differentially expressed genes between sensitive and resistant cells, users can easily perform GO and KEGG enrichment analysis (Fig 3e) via web interface. Enriched pathways are displayed using a bubble plot, where the size represents the number of associated genes and the color encodes statistical significance.
- Cell-cell communication: User can click to calculate and display cell-cell communications between different types of cells using CellphoneDB tool [21]. Specifically, users are guided to focus on the strength of ligand-receptor interactions between resistant and sensitive cells (Fig 3f).
- scRNA-seq data visualization: UMAP is used to display single-cell sequencing data, with cell types annotated using the SingleR tool [22]. Here users can browse the proportion and number of sensitive and resistant cells within a dataset (Fig 3g). Additionally, the expression profiles of genes of interest in different cells can be displayed through a heatmap on the Dataset Detail Page.
- Dataset download: Users can download datasets of interest from the Dataset detail page. More importantly, we have constructed a batch of combined datasets regarding specific drugs, by using the Harmony tool [23] to remove batch effects. These combined datasets are ready for the usage in training deep learning models for predicting single-cell drug response.

**Fig. 3.**
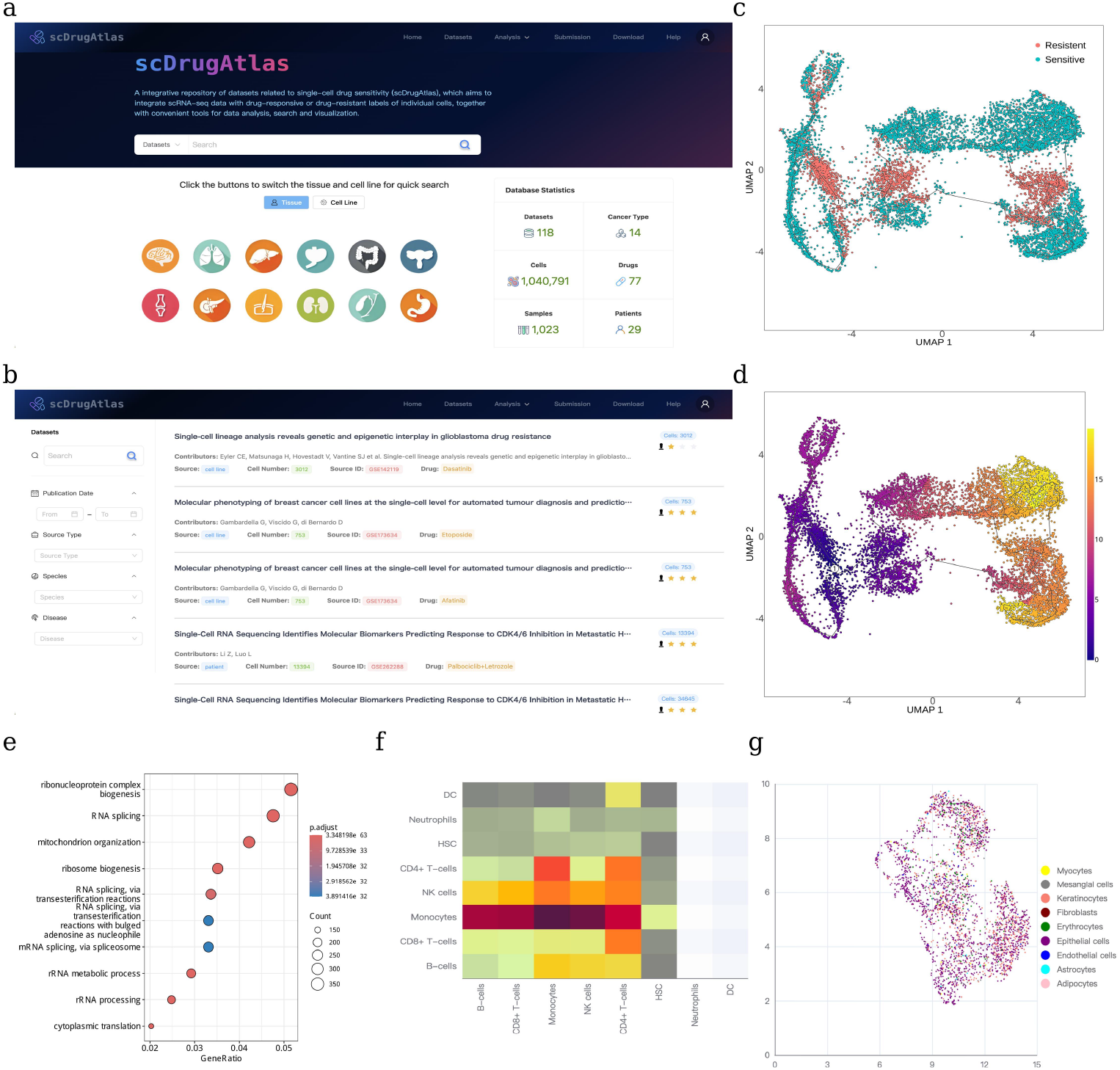
Overview of the scDrugAtlas web platform and its functional modules. (a) Homepage of scDrugAtlas. (b) Dataset listing and filtering interface, allowing users to search by source type, species, release date, and disease. (c) Evolutionary trajectory of tumor cells colored by drug response labels (resistant vs sensitive). (d) Evolutionary trajectory of tumor cells colored by inferred pseudotime values. (e) Functional enrichment analysis based on differentially expressed genes between sensitive and resistant cells. (f) Cell-cell communication analysis between different types of cells. (g) UMAP visualization of scRNA-seq data with annotated cell types.

## 5 Case studies

### 5.1 Cross-study validation on Erlotinib-treated PC9 cells

To evaluate the effectiveness of our curated datasets in building predictive models, we present two case studies. First, we utilized a well-documented scRNA-seq dataset derived from the non-small cell lung cancer (NSCLC) PC9 cell line (GEO accession: GSE196018 [24], scDrugAtlas ID: 126) as the training set. This dataset comprises 9,546 Erlotinib-sensitive cells (labeled as 1) and 9,905 Erlotinib-resistant cells (labeled as 0), all of which exhibit well-defined phenotypic characteristics. Meanwhile, we incorporated another independent scRNA-seq dataset of Erlotinib-treated PC9 cells (GEO accession: GSE149383 [25], scDrugAtlas ID: 113) as test set. This dataset includes individual cells collected at multiple time points following drug exposure. To establish a test set for predictive model, we labeled 765 cells from day 0 (prior to drug exposure) as sensitive cells, while a total of 371 cells from days 9 and 11 post-treatment as resistant ones. The Harmony tool was used to mitigate batch effects between the training and test datasets. Pseudotime analysis of the test set demonstrated that PC9 cells progressively shifted toward drug resistance with prolonged Erlotinib exposure (Fig 4a-b). Next, several typical classifiers—including random forest (RF), logistic regression (LR), support vector classifier (SVC), multilayer perceptron (MLP), and Gaussian naive Bayes (Gaussian NB)—were trained on the training data and subsequently applied to the test data to evaluate their performance. The results showed that RF classifier demonstrated strong predictive power, achieving an accuracy of 87.41% and AUROC of 0.978 (Fig 4c). This cross-study validation provides compelling evidence for the reliability of our dataset and assigned labels in building predictive models.

**Fig. 4.**
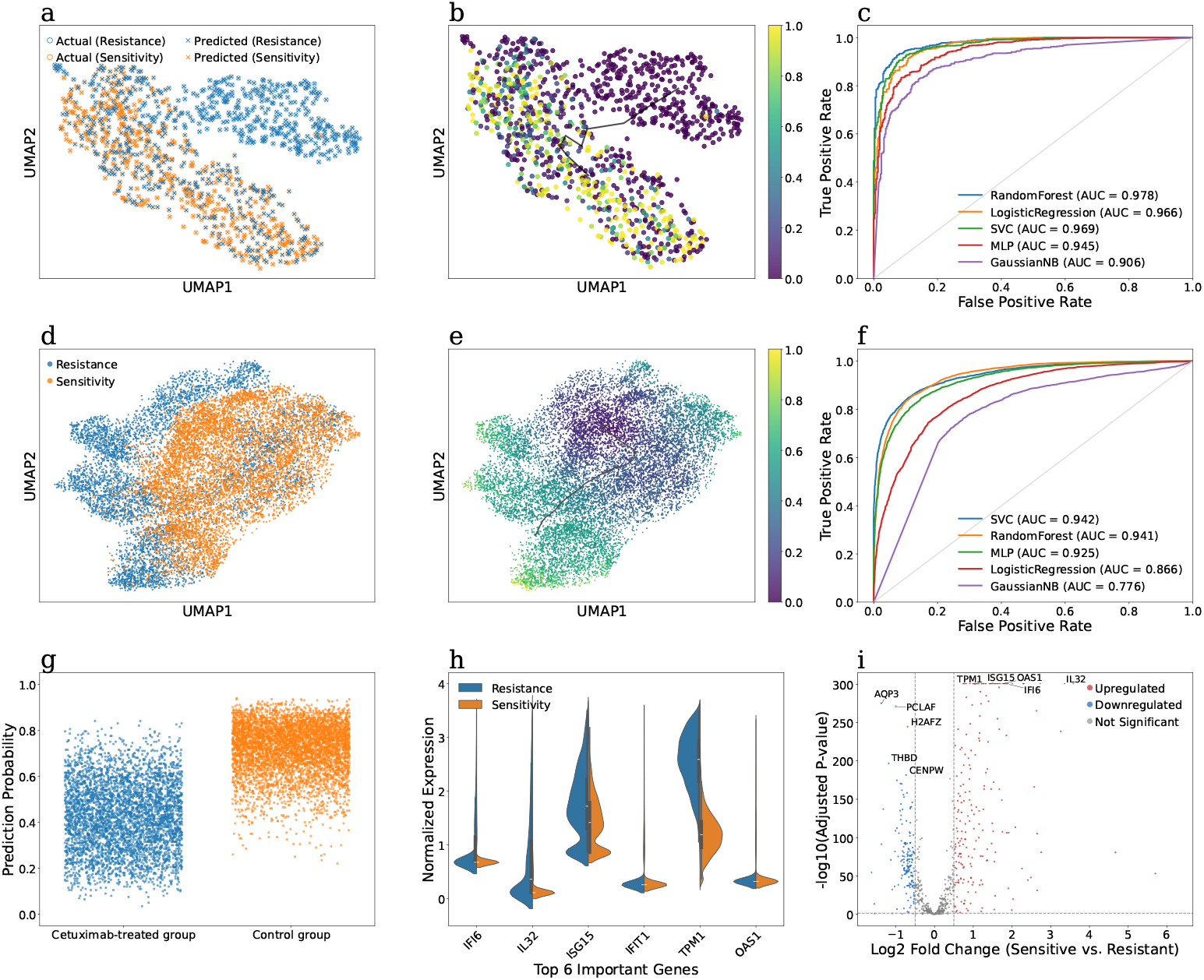
Two case studies demonstrating the development of predictive models for identifying drug-resistant tumor cell subpopulations. Case study 1: (a) UMAP visualization of scRNA-seq data from PC9 cells colored by resistance (blue) and sensitivity (orange) labels, showing the true labels (circle) and the RF-predicted labels (cross). (b) Pseudotime analysis of PC9 cells colored by RF-predicted sensitivity probability (yellow for sensitive; purple for resistant). (c) ROC curves and AUROC values of multiple classifiers evaluated on test set. Case study 2: (d) UMAP visualization of the scRNA-seq data from HNSCC cells, colored by resistance (blue) and sensitivity (orange) labels. (e) Pseudotime analysis of the HNSCC cells colored by the SVC-predicted sensitivity probability (yellow for sensitive; purple for resistant). (f) ROC curves and AUROC values of multiple classifiers evaluated on the test set. (g) Scatter plot of the true labels and SVC-predicted probabilities of individual cells in the test set. (h) Violin plot showing the normalized expression levels of the six most important genes in classifying resistant and sensitive cells. (i) Volcano plot of differentially expressed genes between resistant and sensitive cells.

### 5.2 Cross-cell type validation on Cetuximab-treated HNSCC cells

We further conducted another cross-cell type case study using a Cetuximab-treated head and neck squamous cell carcinoma (HNSCC) dataset comprising 23,914 HNSCC cells from three distinct HNSCC cell lines: SCC1, SCC6, and SCC25 [26]. Among these, 13,040 cells were classified as sensitive (labeled as 1) and 10,874 cells as resistant (label as 0). Integrative analysis of single-cell drug response distribution and pseudotime trajectories revealed that HNSCC cells progressively transition toward a Cetuximab-resistant state (Fig 4d-e).

To build a predictive model across cell types, SCC1 and SCC25 cells were used as training set, while SCC6 was reserved as test set. Similarly, multiple classifiers were trained on the training set and evaluated on the test set. As shown in Fig 4f, SVC achieved the best performance with an AUROC value of 0.942, while both RF and MLP also yielded more than 0.9 AUROC values. These results demonstrated that the predictive models maintained strong generalization ability when applied to the previously unseen SCC6 cell line. To further evaluate the model’s performance at the single-cell level, we examined the predicted probabilities of individual cells in control and Cetuximab-treated groups (Fig 4g). The majority of control cells were assigned high predicted values corresponding to sensitivity, whereas most Cetuximab-treated cells were assigned notably low predicted values indicative of resistance, leading to a clear separation between the two groups. Beyond predictive accuracy, we also explored the key biological features underlying the model’s discrimination between sensitive versus resistant phenotypes. Feature importance analysis identified the most contributive genes, and the expression patterns of the top six (IFI6, IL32, ISG15, IFIT1, TPM1, and OAS1) are shown in Fig.4h. Notably, genes associated with interferon-stimulated pathways, such as ISG15, were significantly up-regulated in resistant cells compared to sensitive cells, consistent with previous studies that have validated their potential roles in drug resistance [27, 28]. Also, up-regulation of IFIT1 and tumor-associated neutrophils has been shown to promote immune suppression and resistance to immunotherapy in poorly cohesive carcinoma[29]. Moreover, differential expression analysis further confirmed that TPM1 and OAS2 are strongly up-regulated in resistant cells (Fig 4i), exhibiting both large fold changes (Log2) and high statistical significance (-log10 adjusted P-value).

Taken together, these case studies demonstrate that our curated datasets not only enable the construction of highly accurate predictive models for identifying drug-resistant cell subpopulations, but also uncover biologically meaningful resistance biomarkers and regulatory pathways, underscoring the substantial potential and practical value of our resource in advancing cancer research.

## 6 Discussion and Conclusion

The phenotype-based high-throughput screening (HTS) primarily captures the overall drug response of tumor cells but falls short in revealing the difference of drug response at single-cell level. This limitation arises because bulk RNA sequencing (bulk RNA-seq) often produces signals dominated by growth-advantageous cell clones, thereby obscuring signals of the minority of drug-resistant cell subpopulations. As a result, the molecular mechanisms underlying tumor drug resistance remain largely uncharacterized.

We also noted other databases that aggregate scRNA-seq data to investigate tumor drug sensitivity and resistance. For instance, scDrugAct [30] offers a large-scale resource integrating single-cell transcriptomic profiles with drug perturbation data to delineate drug–gene–cell associations, highlight tumor microenvironment cell heterogeneity and drug response. Similarly, CeDR Atlas [16] integrates scRNA-seq datasets with CMap drug signatures to infer cell type-specific drug responses, with particular attention to potential side effects on normal cells. While both resources incorporate extensive single-cell transcriptomic data, they rely on drug perturbation signatures from CMap as references for response inference. Consequently, they lack direct singlecell–level evidence of drug response and cannot provide labeled datasets supportive for predictive model training. In contrast, scDrugAtlas places greater emphasis on on building high-quality single-cell drug response datasets, wherein drug response labels are derived from experimental metadata and meticulously curated through manual annotation. Moreover, scDrugAtlas implements a confidence-level grading scheme for the labels based on the experimental evidence. To our knowledge, scDrugAtlas is currently the only database offering manually curated, single-cell–level drug response labels alongside batch-effect–corrected integrated datasets that preserve explicit sensitive/resistant annotations. These features render the resource particularly well suited for training and validating predictive models of single-cell drug response, offering unique advantages in identifying resistant subpopulations and elucidating mechanisms underlying drug resistance. Notably, scDrugAtlas has already demonstrated its utility through two case studies on Erlotinib (NSCLC PC9 cells) and Cetuximab (HNSCC cells), underscoring the value of these curated data resource. Despite the DRMref [17] database also paid attention to the single-cell drug resistance, the scale of data and the diversity of covered drugs and species remain significantly limited. Compared to DRMref, our database possessed nearly three times the amount of data, as well as more reliable drug response labels. To ensure the reliability of the drug response labels, we meticulously verified several factors: the origin of the cell tissue (whether from primary, recurrent, or metastatic lesions), the duration of drug exposure, and the presence of acquired resistance and continued cell proliferation.

In addition to the advantages in dataset volume and label quality, we have also prioritized the development of functionality for comparative analysis between drug-sensitive and drug-resistant cells. By correlating the tumor cell evolutionary trajectories with the progression of cells towards drug-resistant state, users can easily identify the critical state of cellular transition from sensitivity to drug resistance during pseudo-time, which greatly facilitate the discovery of new drug resistance genes. Furthermore, we calculate the activity levels of signaling pathways in drug-sensitive and drug-resistant cell subpopulations, and conduct differential analysis based on the activity levels to reveal the drug resistance mechanisms of tumors at the single-cell level.

